# ciftify: A framework for surface-based analysis of legacy MR acquisitions

**DOI:** 10.1101/484428

**Authors:** Erin W. Dickie, Alan Anticevic, Dawn E. Smith, Timothy S. Coalson, Mathuvanthi Manogaran, Navona Calarco, Joseph D. Viviano, Matthew F. Glasser, David C. Van Essen, Aristotle N. Voineskos

## Abstract

The preprocessing pipelines of the Human Connectome Project (HCP) were made publicly available for the neuroimaging community to apply the HCP analytic approach to data from non-HCP sources. The HCP analytic approach is surface-based for the cerebral cortex, uses the CIFTI “grayordinate” file format, provides greater statistical sensitivity than traditional volume-based analysis approaches, and allows for a more neuroanatomically-faithful representation of data. However, the HCP pipelines require the acquisition of specific images (namely T2w and field map) that historically have often not been acquired. Massive amounts of this ‘legacy’ data could benefit from the adoption of HCP-style methods. However, there is currently no published framework, to our knowledge, for adapting HCP preprocessing to “legacy” data. Here we present the ciftify project, a parsimonious analytic framework for adapting key modules from the HCP pipeline into existing structural workflows using FreeSurfer’s recon_all structural and existing functional preprocessing workflows. Within this framework, any functional dataset with an accompanying (i.e. T1w) anatomical data can be analyzed in CIFTI format. To simplify usage for new data, the workflow has been bundled with fMRIPrep following the BIDS-app framework. Finally, we present the package and comment on future neuroinformatics advances that may accelerate the movement to a CIFTI-based grayordinate framework.

**HIGHLIGHTS:**

- the ciftify package allows for grayordinate-based (CIFTI format) analysis of non-Human Connectome Project (i.e. legacy) MR acquisitions
- The workflow and dependencies are distributed as a Docker container, following the BIDS-app interface
- Additional ciftify utilities aid in downstream analysis of CIFTI images
- We intend for this work to offer bridging solution for legacy data that will allow many researchers to adopt CIFTI format analyses

## 1.0 Introduction

The Human Connectome Project (HCP) has incorporated major technical advances at many steps of neuroimaging data acquisition and analysis (Glasser et al., 2016b; Van Essen et al., 2013). At the level of MR acquisition, the HCP used multi-band MR pulse sequences, which increased both the temporal and spatial resolution of MR data (Uğurbil et al., 2013; Van Essen et al., 2012). In addition, the HCP project took great care to utilize state-of-the-art approaches to correct for MR field bias and image distortions apparent across MR modalities (Glasser et al., 2016b). At the level of MR analysis, as part of the minimal preprocessing pipelines, the HCP introduced a novel Connectivity Informatics Technology Initiative file format (CIFTI; https://www.nitrc.org/projects/cifti/) for conducting analyses in the “grayordinate” framework. In CIFTI format, data from cerebral cortical gray matter is stored in relation to 2-dimensional surface meshes, whereas subcortical data is maintained in 3-dimensions, within the same file, by representing only subcortical gray matter voxels. The HCP also introduced a powerful visualization tool, Connectome Workbench, with an accompanying suite of command-line functions that allow for the manipulation of CIFTI, GIFTI and NIFTI format images (Marcus et al., 2013).

A 2D cortical surface-based approach to MR analysis of cortical signals provides several advantages over the more commonly used 3D volume-based approach as previously described (Argall et al., 2006; Fischl, 2012; Goebel et al., 2006; Zijdenbos et al., 2002). Most notable benefits include better adherence to the inherent geometry of cortical surfaces, increased statistical power (Anticevic et al., 2008; Argall et al., 2006; Coalson et al., 2018; Fischl et al., 1999; Jo et al., 2007; Tucholka et al., 2012), removal of the deleterious effects of volume-based smoothing (which markedly erodes spatial localization (Coalson et al., 2018), superior visualization (Van Essen, 2012), and finally a simplified, more compact framework for multimodal analysis (such as analyses combining surface-based anatomical features, e.g., cortical thickness, and fMRI). If should be noted that these benefits have been reported using data with lower spatial and temporal resolutions than the HCP. For these reasons, surface-based analyses have provided new insights into how the human cortex is functionally organized within humans at both the population level (Glasser et al., 2016a; Margulies et al., 2016; Mueller et al., 2013; Van Essen and Glasser, 2018; Yeo et al., 2011) and individual level (Braga and Buckner, 2017; Glasser et al., 2016a; Wang et al., 2015).

Although the benefits of surface-based over volume-based registration have been widely recognized for nearly a decade and despite the continued development and promulgation of surface-based tools by several groups (Argall et al., 2006; Fischl, 2012; Goebel et al., 2006; Van Essen, 2012; Zijdenbos et al., 2002), the adoption of surface-based analyses by the neuroimaging community has been slow. One reason for this inertia is that many widely used tools for MR preprocessing (e.g., SPM, FSL) as well as widely adopted pipeline tools (e.g., NIAK, CPAC) do not offer “out of the box” workflows with surface-based registration steps. These tools are not designed to support the CIFTI format, or provide visualization frameworks for combined 2D/3D geometry. The HCP consortium released its pipelines as part of the Minimal Preprocessing Pipeline GitHub project (Glasser et al., 2013). One key requirement of the Minimal Preprocessing Pipeline is acquisition of a high-resolution T2-weighted image, used to generate high-quality surface reconstructions (Glasser et al., 2013) and myelin maps (Glasser et al., 2014; Glasser and Van Essen, 2011), that was not typically acquired in legacy human MR protocols. Therefore, many ‘legacy’ acquisitions often collected without high-resolution T2 images or fieldmaps cannot be processed using the HCP’s pipelines.

It is estimated that tens of thousands of participants are scanned annually as part of research studies, and this number has been growing for over 25 years (Smith, 2012). Thanks to data sharing consortia such as the International Neuroimaging Data-Sharing Initiative (INDI, (Milham et al., 2018)) and OpenfMRI (Poldrack et al., 2013), thousands of legacy datasets are publicly available (Eickhoff et al., 2016; Poldrack and Gorgolewski, 2014). The National Institute of Mental Health (NIMH) have also committed to data sharing via the NIMH Data Archive (NDA; ndar.nih.gov). These legacy datasets are drawn from healthy individuals plus a wide variety of clinical populations and developmental stages. While some clinical and developmental populations are now being scanned according to HCP acquisition standards, it may be years before these sample sizes will be large enough to answer some of today’s most pressing questions. The HCP requirements can be viewed either as a barrier or as an opportunity. If viewed as an opportunity, there is a concomitant need to develop tools for HCP-style analyses applicable to legacy MR data available today (and likely the near future) Therefore, maximally leveraging large legacy datasets is important for clinical research, especially for those attempting to characterize disease heterogeneity (Choudhury et al., 2014).

Here, we address the opportunity to leverage important innovations of the HCP pipelines to enable integration of two decades of existing legacy human neuroimaging data into the CIFTI grayordinate-based framework. To expand the utility of HCP-style methods, we present the ciftify package for grayordinate-based (CIFTI format) analysis of legacy acquisitions that have already been processed using FreeSurfer. (Datasets that use other cortical segmentation methods or have no surface-based processing at all are considered in the discussion.) Ciftify translates two key modules of the HCP Minimal Preprocessing Pipeline: the FreeSurfer-to-Connectome Workbench conversion, and the fMRI surface projection, into simple command line tools. Integrating these two steps will work from FreeSurfer-based anatomical outputs (generated from only T1w images) to convert current volume-based fMRI analysis pipelines into a grayordinate-based one. These tools allow researchers to move their analyses to the surface while ensuring the opportunity to analyze non-HCP quality data. Below, we describe the ciftify package and discuss its expected use case. We also introduce additional tools for running and interpreting group-level analyses in CIFTI format.

## 2.0 The ciftify preprocessing workflow and BIDS-app

A diagram of the ciftify preprocessing workflow is given in <u>Figure 1</u>. As a precursor to ciftify, surfaces are generated from T1w anatomical images using FreeSurfer’s recon_all function (Fischl, 2012), and fMRI runs are preprocessed using other software as discussed below. The BIDS-app Docker container will use FMRIPrep (Esteban et al., 2019a, 2019b) for these preprocessing steps if they have not already been run. Anatomical data is converted from FreeSurfer to CIFTI formats, and MNI inter-subject anatomy-based registration and resampling is performed by the ciftify_recon_all function (using FSL’s FNIRT). Next, fMRI acquisitions are projected to the surface, and subcortical data are resampled by the cifitfy_subject_fmri function. Quality assurance visualizations for these steps are generated with the cifti_vis_recon_all and cifti_vis_fmri utilities.

**Figure 1.**
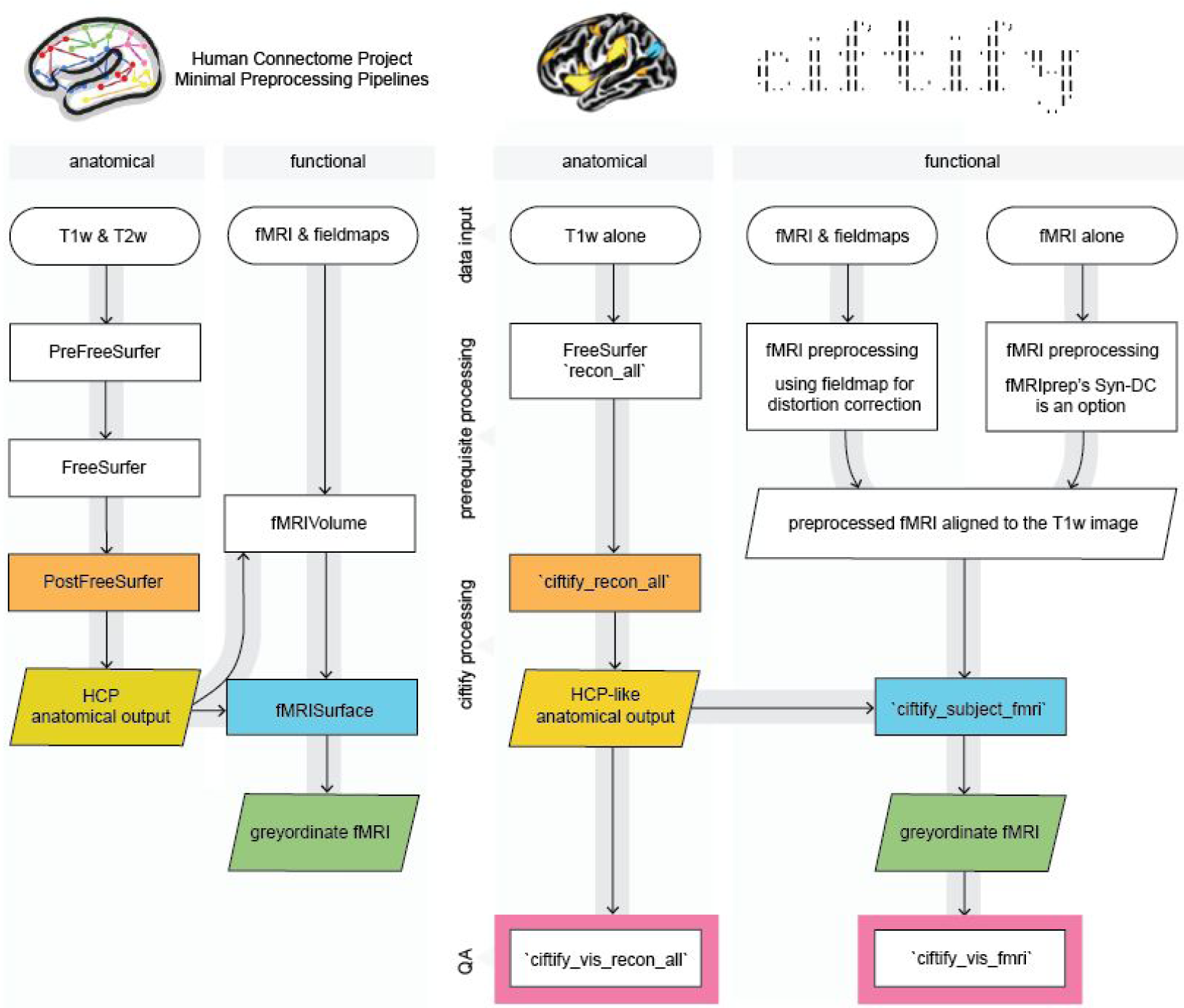
Key modules from HCP minimal processing pipelines (left) were adapted to create the ciftify preprocessing workflow (right). Within the fmriprep_ciftify BIDSapp the prerequisite processing steps listed here are performed by fmriprep. Additional usage options, including the ability of the user to input pre-calculated transforms to standard space, are given at https://edickie.github.io/ciftify/.

### 2.1 Step 0: Prerequisite processing to ciftify

The ciftify approach is achieved starting from the standard FreeSurfer recon-all output derived from at least one raw T1w image. The requirements for the T1w image is that is can be used to produce freesurfer surface outputs passing quality assurance. Therefore, like FreeSurfer, we recommend a resolution no greater than 1mm isotropic. The ciftify approach to fMRI should, in theory, work on any Freesurfer output along with any fMRI volume that has been preprocessed with any fMRI preprocessing pipeline, assuming that 1) no volume-based smoothing has been applied and 2) the image has not been non-rigidly warped into standard space (Note: optional, expert user options allow for a standard space functional image to be provided with additional files detailing the standard space warp performed, see online documentation at https://edickie.github.io/ciftify/). To ensure the quality of the surface mapping step, it is recommended that preprocessing prior to ciftify_subject_fmri include steps to correct for EPI signal distortions. The fMRIPrep pipeline (Esteban et al., 2019a, 2019b) incorporates many recommended practices for these fMRI preprocessing steps, as well as running the FreeSurfer pipeline (version 6.0). Therefore, the fmriprep_ciftify BIDS-app will run FMRIPrep to accomplish these preprocessing steps. Those wishing to run ciftify without the FMRIPrep preprocessing base can run the individual ciftify steps manually. More instructions for doing so are given at https://edickie.github.io/ciftify/#/tutorials/example-usage. Recommendations for EPI distortion correction step are detailed further in *4.2 Functional preprocessing considerations*.

Importantly, the quality of surface-mapped MRI is strongly influenced by the registration between the functional volume and the anatomical data, which suffers in areas where the EPI image is distorted. HCP pipelines include two preprocessing steps to maximize the quality of registration across MR modalities: the acquisition and use of b0 field inhomogeneity maps (Glasser et al., 2013), and the use of Boundary-Based Registration (BBR) for cross-modal alignment (Greve and Fischl, 2009). The FMRIPrep pipeline is a “glass box” workflow that allows for a simplified user interface for the implementation of these recommended approaches for fieldmap-based EPI distortion correction and Boundary-Based Registration (BBR) for cross-modal alignment. In addition, FMRIPrep offers an option for “fieldmap-less” distortion correction, which estimates the areas of distortion using a combination of a participant’s T1w anatomical data and an average fieldmap template (Treiber et al., 2016; Wang et al., 2017).

### 2.2 Step 1: The participant anatomical workflow

The first step in the ciftify workflow is accomplished by `ciftify_recon_all,` a command line utility adapted from the PostFreeSurferPipeline module of HCP’s Minimal Preprocessing Pipeline. Outputs from FreeSurfer are converted to GIFTI and CIFTI format. In turn, surface-based alignment of the cortical mesh is performed using the MSMSulc algorithm (Robinson et al., 2018) followed by resampling to a 32k standard. In line with the HCP’s Minimal Preprocessing Pipeline, `ciftify_recon_all` outputs a directory structure where output files are divided into subfolders representing analysis “spaces”. More detail on these “spaces” is given in Glasser and colleagues (2013). In general, the MNINonLinear/fsLR32k combines the benefits of MNI volumetric registration (for subcortical structures) with surface-level registration based on sulcal anatomy for the cortex. This 91,282 standard-mesh “grayordinates” space, resampled to an approximate ‘32k’ (average ∼2mm spacing), is the HCP’s standard space for human fMRI and multimodal analyses (note that HCP fMRI was acquired with 2mm voxel resolution). The surface meshes for fsLR32k space differ from the fsavarge meshes (used in FreeSurfer pipelines) not only in resolution (fsaverage5 has ∼4mm spacing), but also (and more importantly) by providing mirror-symmetry between the left and right meshes (i.e., vertex 1024 on the left mesh corresponds in geographic location to vertex 1024 in the right mesh (Van Essen, 2012). This mirroring facilitates the analysis of symmetries and asymmetries in brain connectivity and diverse cortical features. Importantly, the CIFTI format also includes the FreeSurfer volumetric segmentation of subcortical and cerebellar structures.

For the purpose of retaining data provenance, `ciftify_recon_all` also produces a log file that records information about the input arguments, software environment, and information copied from the FreeSurfer logs about the environment in which recon_all was run. These logs also record all executed command line calls, so that the user can review the intermediate steps performed.

Due to the widespread use of FreeSurfer in the scientific community, it is anticipated that many investigators may already have recon_all outputs generated for their specific projects, with appropriate quality assurance completed. For such studies, the process to convert this data to CIFTI format can be achieved relatively quickly. However, if a high-resolution T2w image is available (i.e. a T2w image of voxel size matched to the T1w image), it is recommended that the HCPPipelines anatomical modules are run (PreFreeSurfer, FreeSurfer, PostFreeSurfer) instead of FreeSurfer and cifitify_recon_all (see *4.1 Anatomical preprocessing considerations*).

The anatomical workflow also produces quality assurance (QA) images, using the `cifti_vis_recon_all` utility, for reconstructed surfaces and subcortical masks (<u>see Figure 2A</u>). These images are analogous to those produced for other FreeSurfer quality assurance frameworks (such as FreeSurfer’s own QATools). Some investigators may find these outputs to be a valuable adjunct to FreeSurfer recon_all outputs. An optional flag will output an accompanying “.scene” file, which allows the user to browse these QA views interactively in the connectome-workbench viewer. These QA views include slice representations with pial and white surface contour overlays, as well as the reconstructed surfaces (derived from the midthickness surface) with FreeSurfer’s automatic parcellation (aparc) anatomical labels viewed from multiple standard angles. These images are generated using both the native and MNI space version of the outputs. Also, slices showing the masks generated by the FreeSurfer Automatic Segmentation (aseg) are presented in native T1w space so that these masks can be evaluated.

**Figure 2.**
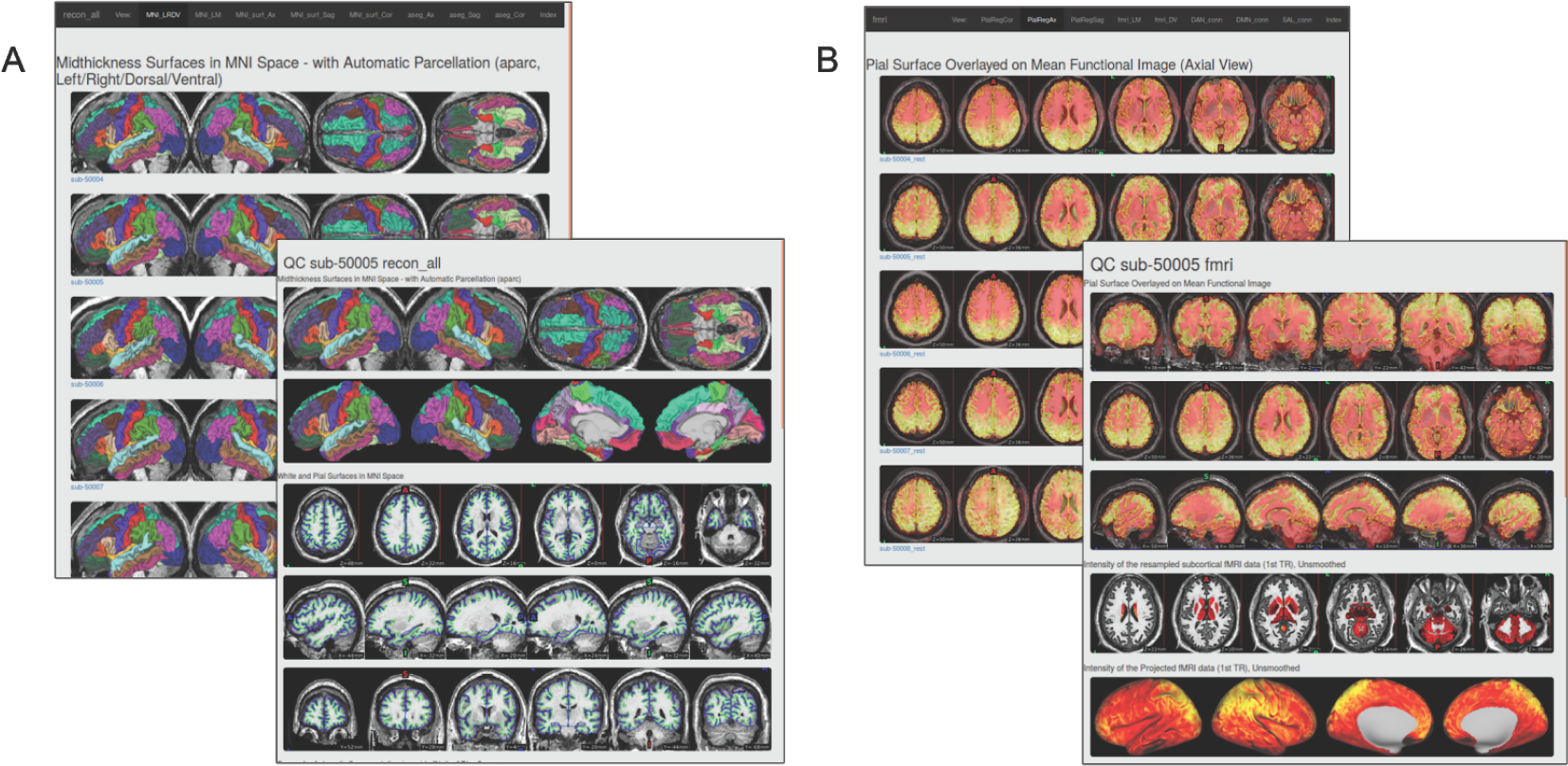
Example quality assurance (QA) pages created using A) cifti_vis_recon_all and B) cifti_vis_fmri. These example QA webpages can be viewed at https://edickie.github.io/ciftify/demo/qc_recon_all/index.html.

### 2.3 Step 2: The participant functional workflow

Once surface anatomy has been defined for a particular subject, the `ciftify_subject_fmri` utility is employed. The ciftify_subject_fmri stage maps preprocessed fMRI volumes to a specific subject’s fsLR23k “grayordinate” space. This utility, adapted from the fMRIsurface module of HCP’s Minimal Preprocessing Pipeline, projects from volume to the surface in a weighted, ribbon-constrained manner, where the value assigned to each vertex is calculated as a weighted average of the voxels encompassed (or partially encompassed) by the cortical ribbon. Excessively noisy voxels, defined according to their local coefficient of variation, are excluded from the mapping. Importantly, this surface mapping approach differs from those used in other known pipelines - such as FreeSurfer, that resample only from voxels that intersect with the midthickness surface. Another notable difference from previous approaches is that data from subcortical structures are resampled within their anatomical masks to approximately 32k subcortical voxels (including cerebellum). The final output is a CIFTI mapped (dtseries.nii) version of the functional data, in MNINonLinear, fsLR32k space. Note that the fMRI input volume should be “minimally” preprocessed with no volume smoothing and no non-rigid registration to standard space (else that same registration would need to be applied to the surfaces too, which is not supported by ciftify). From this step, optional additional CIFTI smoothing can be added (2D surface smoothing for the cortical ribbon and parcel-constrained smoothing for subcortical volumes), though see the cautionary note in Coalson et al (2018) about smoothing of any kind.

Like the anatomical workflow, the functional workflow also produces visualizations for QA of the volume-to-cortex mapping step as well as subcortical resampling (<u>see Figure 2</u>B), using the cifti_vis_fmri utility. These images include slice overlays of the functional volume with the pial surface and views of the unsmoothed functional signal on the surface.

### 2.4 Generating a group quality assurance interface

These images are then combined into HTML pages so that collections of images can be viewed together in a browser window. Importantly, the HTML pages present the QA images in both subject-level pages (where all images related to one scanning session are presented together) as well as population-level index pages. These pages present the same view for all subjects in a study together. When the population is displayed beside each other, on the same page, QA failures can be identified more easily.

The cifti_vis utilities employ wb_command’s ‘-show-scene’ functionally to generate images for visual inspection. An optional flag will output an accompanying “.scene” file, which allows the user to browse these QA view options interactively in the connectome-workbench viewer. This framework does not require an external display nor MatLab proprietary software, which allows for easier generation on high-performance clusters, thus providing an advantage compared to other QA image-generating tools.

### 2.5 Programming Environment and Usage

The ciftify package contains a set of command-line utilities coded in Python. All code is publicly available under the MIT license at https://github.com/edickie/ciftify. The ciftify Python package (version 2.3.2) can be installed locally using the Python package management system (i.e., pip). Like the HCP pipeline, the ciftify package depends upon publicly available MR analysis software packages (FSL, FreeSurfer and Connectome Workbench) for various conversion and image manipulation steps. Also in concordance with the HCP Pipeline, the default surface registration algorithm is MSMSulc, dependent on the MSM registration tool (https://github.com/ecr05/MSM_HOCR/releases), and the Python packages docopt (docopt.org), nibabel (Brett et al., 2019), nilearn (Abraham et al., 2014), pybids (Yarkoni et al., 2019), NumPy (Walt et al., 2011), SciPy, PyYAML (https://pyyaml.org/), pandas (McKinney, 2010), seaborn (Waskom et al., 2017) and Matplotlib (Caswell et al., 2019) are dependencies. For further usage instructions and example code, see https://github.com/edickie/ciftify/wiki.

This complete programming environment is available as a BIDS app (Gorgolewski et al., 2017) available on Docker Hub (https://hub.docker.com/r/tigrlab/fmriprep_ciftify/). Therefore, the workflow should be relatively straightforward to set up and run for any dataset organized in accordance with the Brain Imaging Data Structure (BIDS; (Gorgolewski et al., 2016).

Example Docker usage:

~~~
docker run -ti --rm \
   -v /path/to/bids/raw/data:/bids_in:ro \
   -v /path/to/output:/derivatives \
   tigrlab/fmriprep_ciftify:latest /bids_in /derivatives participant
~~~

As with all BIDS apps the <bids_in> should be the top level bids directory containing all data participants data for the study. The <derivatives> output directory path will be populated with three subfolders of outputs (freesurfer/ fmriprep/ and ciftify/). If one has already completed part of the processing (i.e. FreeSurfer) these processing steps will be skipped by the BIDS app if the output folders are already present. The “bids_in” input argument provides a path to the dataset to be analyzed (read-only), which must conform to the BIDS standard (see the BIDS speciftication (http://bids.neuroimaging.io) and Supplementary Table 1 for more details of the input file structure). The above command would run the ‘participant’ level workflow for all available functional data in the dataset in a serial fashion. However, additional flags can be added select specific participants, sessions or tasks for processing. These flags are valuable for splitting up participant level processing into smaller tasks to be submitted to a computing cluster.

For example, one could select to only process the resting state task data from session 015 of the MyConnectome data (as in the above figure), using the following command:

~~~
docker run -ti --rm \
  -v /path/to/local/ds000031:/bids_in:ro \
  -v /filepath/to/output:/derivatives \
  tigrlab/fmrirep_ciftify:latest /bids_in /derivatives participant \
  --participant_label=01 \
  --task_label=rest \
  --session_label=015
~~~

Once all participants have been run, a final ‘group’ level mode will quickly write group level (index) quality assurance html pages.

~~~
docker run -ti --rm \
   -v /path/to/data/dir:/data:ro \
   -v /filepath/to/output/dir:/out \
   tigrlab/fmriprep_ciftify:latest /bids_in /derivatives group
~~~

The ciftify Docker container is intended to be run on personal computers and cloud services; however, Docker requires root or root-like permissions, and therefore may not be permitted in many multi-user environments, like High-Performance Clusters, for security reasons. For those using High-Performance Clusters, we recommend using the Singularity software. A Singularity image can either be built directly from the Docker specification (Singularity version >= 2.5), or the docker2singularity converter can be used to convert a Docker image into a Singularity image. For more detailed instructions for each use-case see (https://edickie.github.io/ciftify/#/01_installation).

For example, to build a Singularity image from the DockerHub specification:

~~~
singularity build /my_images/fmriprep_ciftify_image.simg
docker://tigrlab/fmriprep_ciftify:latest
~~~

To run the Singularity image:

~~~
singularity run --cleanenv \
  -B /path/to/bids/raw/data:/bids_in:ro \
  -B /path/to/output:/derivatives \
  fmriprep_ciftify_image.simg /bids_in /derivatives participant
~~~

The output directory structure is in similar to that of the HCP-Pipelines, is organized into folders according to the “space” in which the files are registered. Outputs are comprise of ∼122 files approx 670 MB per participant. (See Supplementary Table 2 for more detailed list of all files generated).

## 3.0 Running analyses in CIFTI format

After data has been preprocessed into CIFTI format, the researcher needs to manipulate CIFTI files, in order to extract measures or run group analyses. For a researcher experienced only in volume-based analysis, this may present a challenge, as code may need to be adapted or rewritten. With recent software developments, most (if not all) calculations previously run in the volume are now possible on the surface. The Connectome Workbench command-line utility (wb_command) offers an extensive suite of functions that are well documented and computationally efficient (Marcus et al., 2013; https://www.humanconnectome.org/software/connectome-workbench). In addition, the nibabel Python package (http://nipy.org/nibabel/) can read and write NIFTI, GIFTI and CIFTI formats, allowing for custom computations in the MatLab or Python (NumPy) environments. Connectome Workbench offers conversion utilities (-cifti-convert) that convert CIFTI data to a “fake” NIFTI (i.e., a file that can be read and written by NIFTI utilities, but that does not contain correct spatial information) so that analyses that have been written for the NIFTI file format can be run. The greatest challenge is presented by those analyses that need to take spatial information into account (e.g., calculations of the spatial extent of a statistical cluster). Fortunately, the command line version of FSL’s statistical packages, such as FSL’s linear modelling tools (FILM and FLAME), MELODIC and FSL’s Permutation Analysis of Linear Models (Winkler et al., 2015, PALM; 2014), for groupwise statistical tests, handles surface data and cifti format.

### 3.1.0 Additional utilities and ciftify binder.org learning environment

The ciftify package also contains several utilities to assist in CIFTI-based surface-level analyses (see Table 1). These utilities are not meant as a replacement for the functionality in Connectome Workbench. In many cases, they are simple utilities that chain together multiple calls to Connectome Workbench. In these cases, commands can be echoed to the user when these utilities are run in with the “--verbose” or “--debug” flags. In some cases (such as the ciftify_meants), data is read into NumPy arrays using the NiBabel package. All code and examples are publicly available on our GitHub repository (https://github.com/edickie/ciftify), which we hope can serve as example workflows for researchers new to the CIFTI analyses.

**Table 1.** Additional utilities included in the ciftify package

| Name | Description |
| --- | --- |
| `ciftify_meants` | Produces a comma separated values (*.csv) file mean voxel/vertex timeseries from a functional file <func> within a seed mask <seed>. <func> functional data input can be NIFTI or CIFTI (*.dscalar.nii or *.dtseries.nii). Seed mask input can be in NIFTI, CIFTI (*.dlabel.nii or *.dscalar.nii) or GIFTI. |
| `ciftify_seed_corr` | Produces a seed correlation map of the mean timeseries within the seed with every vertex/voxel in the functional file. Like `ciftify_meants`, can take a combination of NIFTI, CIFTI or GIFTI inputs. The output seed correlation map matches the file type (NIFTI or CIFTI) of the input functional file. |
| `ciftify_clean_img` | Performs a combination of signal “cleaning” steps (bandpass filtering, confound regression, dummy TR removal and smoothing) for use on CIFTI and NIFTI inputs. Signal cleaning settings can be specified in combination in a “.json” configuration file, or as command line argument. |
| `ciftify_statclust_report` | Creates a tabular statistical report from an input (*.dscalar.nii) map.<br>Includes overlap of clusters with standard atlases |
| `ciftify_atlas_report` | Creates a tabular report of overlap between the input atlas or labels (*.dlabel.nii) with standard atlases |
| `ciftify_vol_result` | Simplified mapping of a NIFTI statistical map or atlas to fsLR32k CIFTI “grayordinate” space of a specified subject. |

In contrast to the full ciftify preprocessing workflow, which requires more than 10 Gigabits of neuroimaging software prerequisites, these tools require only python as well as connectome-workbench. Therefore, they should be easier to install for new learners in a classroom or workshop setting. The programming environment is also available in an online interactive instance on binder.org (https://mybinder.org/v2/gh/edickie/ciftify/master). Future work intends to build interactive examples into this public, online, tutorial space.

### 3.1.1 ciftify_statclust_report

The ciftify_statclust_info utility generates a table (.csv output) listing statistical clusters defined from an input CIFTI (.dscalar.nii) statistical map. A CIFTI file of labelled clusters is also output. The table lists, for each cluster, the cluster ID and name (matching the label in the corresponding dlabel.nii output), as well as the surface area of that cluster (calculated on the midthickness surface). The table also reports the overlap of the cluster with three atlases: (i) the FreeSurfer anatomical atlas (Desikan et al., 2006); (ii) the Yeo seven resting state networks atlas (Yeo et al., 2011); and (iii) the Glasser Multimodal Parcellation atlas (Glasser et al., 2016a). For each peak, for each atlas, the table reports the label in which the peak vertex falls, as well as the proportion of the cluster (from the statistical input map) that overlaps with that index <u>(see Figure 3)</u>.

**Figure 3.**
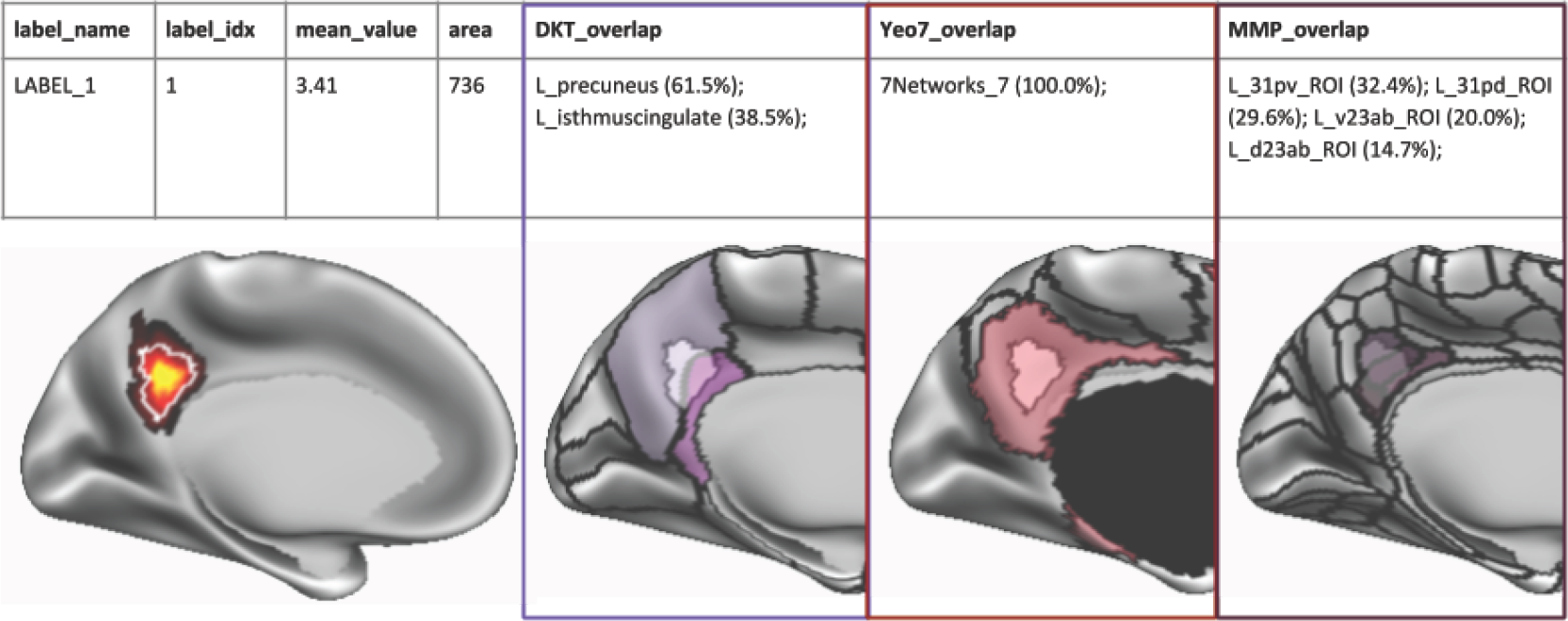
Example output table generated by ciftify_statclust_report. For this simulated statistical map (red-yellow shading), the output table summarizes a statistic map, according to input statistical thresholds, as one cluster (white outline). From the cluster, ID number, mean statistical value and the approximate surface area in mm^2^. Then, for three surface atlases, we report atlas labels encompassing the proportion of the cluster’s surface area that overlaps with this atlas label. In this example, 61.5% of the cluster overlaps the “L_precuneus” label of the FreeSurfer anatomical atlas (Desikan et al., 2006). Likewise, the entire cluster falls within Network 7 (i.e. the Default Mode Network) of the Yeo atlas (Yeo et al., 2011). Lastly, 32.4% of the cluster’s surface area falls within the ‘L_31pv_ROI’ label from Glasser and colleagues Multimodal Parcellation atlas (Glasser et al., 2016a).

## 4.0 DISCUSSION

The brain imaging field has lacked a framework for leveraging the HCP preprocessing philosophy for non-HCP legacy data. The ciftify project presented here helps to close this gap and encourages wider adoption of surface-based analysis of MR data, specifically using the CIFTI file format, for groups working with non-HCP legacy datasets. Allowing researchers to adapt the preprocessing pipelines they are most familiar with allows for 1) easier uptake of surface-based analysis insofar as we remove a steep learning curve and 2) a more flexible framework whereby we can test different preprocessing steps in order to optimize surface-level results across various acquisition settings. For this purpose, the workflow (fmriprep_ciftify), packaged in as a BIDS-app, adapts two critical sections from HCP’s Minimal Preprocessing Pipelines and can be readily deployed on legacy datasets. This workflow also integrates the generation of quality assurance visualizations. In addition, the ciftify package contains a suite of tools to aid in the analysis and manipulation of data after it has been preprocessed into CIFTI format.

### 4.1 Anatomical preprocessing considerations

The goal of ciftify_recon_all was to start directly from default recon-all FreeSurfer outputs. Thus it offers a relatively simple starting point for groups to adopt and test: generating a CIFTI file of surface-level structural measures (such as cortical thickness) is just one command line call away from the workflow that many researchers are already familiar with. While this is an attractive prospect, we feel it important to stress that, when a high-resolution T2w image is available, the additional preprocessing steps incorporated into HCP’s Minimal Preprocessing Pipeline (especially the adapted FreeSurfer approach that uses the T2w signal for better pial surface placement) will yield superior results. To be even more explicit, although the results generated from ciftify will retain many advantages over volume-based processing, they are not equivalent to the results generated from the HCP’s minimal preprocessing pipelines. Note that by “high resolution” T2w image, we are referring to a T2w, or FLAIR acquisition matched in voxel size to the T1w image (i.e. no greater than 1mm isotropic). Hence many existing studies that acquired T2w or FLAIR scans with thick slices or gaps of 2-7mm would not be appropriate. Therefore, the ciftify_recon_all workflow is only recommended for cases when a high-resolution T2w image is not available. Note that the anatomical outputs from the HCP’s Minimal Preprocessing Pipeline can be substituted for ciftify_recon_all outputs for those who would like to use ciftify’s other tools (such as ciftify_subject_fmri).

There remains an ongoing debate regarding the best algorithms, parameters and software for accurate between-subject alignment. We argue that the best registration algorithm should achieve the best alignment for a given level of distortion, or the least distortion for a given level of alignment. This concept applies to both volume-based and surface-based alignment steps within the ciftify pipeline. For the volume-based transformation to standard (i.e. MNI) space, we prefer FNIRT. Compared to the ANTs implementation in fmriprep, FNIRT is tuned conservatively to generate less distortion and as a result, does not overfit to folding patterns <u>(Coalson et al. 2018 see Supplementary Figures S11 and S12)</u>. This recommendation for volume-based alignment mirrors recent observations for surface-based alignment, where the FreeSurfer default alignment produces more spherical distortion and performs less well at aligning cortical areas than does MSM alignment that has been tuned for less distortion (the amount of distortion allowed was determined by optimizing for maximal task-based functional alignment <u>(Robinson et al. 2018; Robinson et al. 2014)</u>. However, as both volume and surface alignment tools continue to be developed, mechanisms need to be available for available tools to be validated in comparison to one another, within the context of pipelines like ciftify. Therefore, we have added expert-user options to change realignment settings or to override the realignment with a user-provided (FSL format) transform file.

### 4.2 Functional preprocessing considerations

Importantly, the quality of surface-mapped MRI is a product of the quality of the volume-to-surface mapping step. This, in turn, is a direct consequence of the registration between the functional volume and the anatomical data, which suffers in areas where the EPI image is distorted. The HCP adopted state of the art methods to maximize the quality of registration across MR modalities. These methods included upgrading sequences to acquire data at a voxel size smaller than average cortical thickness, the acquisition of data for correcting EPI distortion (Glasser et al., 2013), and the use of Boundary-Based Registration (BBR) for cross-modal alignment (Greve and Fischl, 2009). While it is clear that legacy acquisitions will generally not match the quality of HCP, it is important that investigators make preprocessing choices that will maximize the data quality of their dataset given its limitations. A critical step for good fMRI-to-anatomical mapping is correcting for fMRI distortions. Correcting for susceptibility distortions in fMRI data can be done with several approaches. The preferred technique is to use data from additional field map or phase reversed spin echo scans. Phase encoding polarity techniques combine images with opposing phase encoding to calculate distortion maps (using tools such as FSL’s TOPUP (Andersson et al., 2003). This is the technique currently used by the HCP. Second, field maps can be estimated using sequences that measure phase evolution in time between two close gradient echo acquisitions (Hutton et al., 2002). It is becoming common practice to incorporate one of these techniques into current acquisition protocols. However, proper incorporation of these sequences into a fMRI preprocessing pipeline can be challenging for a novice brain scientist, as they often require increased knowledge of the specific scan acquisition settings. The fMRIprep pipeline offers a helpful interface in this regard, where the appropriate EPI distortion software will be called as long as the input files are organized in accordance with the Brain Imaging Data Structure (BIDS). For older acquisitions, where no field map or reverse-phase encoded maps are available, FMRIPrep has implemented a displacement field estimation via nonlinear registration. Another experimental approach is the incorporation of mean field map images (if they are available) from the employed scanner. Future work will explore these and other distortion correction options in legacy fMRI data to ensure that the deformations they generate lead to net improved registrations over no correction.

Another important consideration is whether or not to smooth data on the surface or within the subcortical grey matter structures. Spatial smoothing increases signal to noise ratio, but at a cost to anatomical specificity. It is extremely important to avoid volume-based smoothing before the volume-to-surface mapping step because such smoothing (even with a small 4mm FWHM kernel) is very deleterious to cortical anatomical specificity (Coalson et al., 2018). Once fMRI data is in cifti format, 2D surface-based smoothing can be applied to the cortical data and 3D parcel-constrained smoothing can be applied to the subcortical data. If smoothing is used, we recommend moderation, as even 4mm FWHM in the surface is mildly deleterious, particularly to cortical areas that are narrow (Coalson et al., 2018), but may provide an acceptable tradeoff to increase signal to noise ratio in small sample sizes (in large datasets like the HCP smoothing is often unnecessary). A preferable alternative for improving the signal to noise ratio are approaches like parcellation (Glasser et al., 2016a) or using Wishart-based filtering to preferentially smooth noise over spatiotemporally structured signal (Glasser et al., 2016b). Parcellation is especially preferable to smoothing for researchers who are interested in effects at the level of cortical areas.

### 4.3 The value of higher resolution data

While we advocate use of the HCP analytic approach to all MR datasets, we recognize the synergistic benefits from combining the HCP’s analytic approach with the HCP’s higher standards for MR data acquisition. In particular, higher spatial resolution for MR data, specifically a voxel size less than the mean cortical thickness (∼2.6mm) allows for better separation of signal across sulci/gyri of opposite grey matter ribbons (Glasser et al., 2016b). That said, even fMRI resolution as low as 4mm benefits from the HCP-Style approach to cross-subject alignment and not smoothing in the volume (Coalson et al., 2018). In addition, increasing the temporal resolution of MR data allows for more effective cleaning for MR artifacts from motion or physiology (Glasser et al., 2017, 2016b; Salimi-Khorshidi et al., 2014).

### 4.4 A note about visualizing a volume-based result on the cortical surface

It is becoming more common for researchers to provide surface visualizations when reporting the results of a volume-based group analysis. This practice is beneficial, in comparison to slice representations, for the communication of whole brain results, and can drive more thoughtful interpretations that reflect large-scale networks. For this reason, ciftify_vol_result, a utility for doing simple NIFTI to CIFTI mapping, is included in the ciftify package. If a group average ‘subject’ (i.e. HCP_S1200_GroupAvg) is specified, ‘average fiducial mapping’, or mapping to average surfaces, is performed (Van Essen, 2005). This type of mapping, while fast and easy, suffers from the drawback that group average surfaces fail to closely overlap with the actual location of the cortical ribbon. Instead, they “drift” towards the inside of curvature (often towards white matter) especially in areas of variable folding patterns, such as the association cortex (Coalson et al., 2018). What is more troubling is that if the correspondence between MNI coordinates and cortical areas is not uniform throughout the cortex, results will be most severely impacted in regions of high individual variability and especially for gyral crowns and sulcal fundi. An alternative approach is to repeat the volume-to-surface mapping steps using the surfaces from a collection of subjects, and then to summarise (i.e., average) the results. This so-called “multi-fiducial mapping” (Van Essen, 2005) samples more uniformly from association cortex, with the caveat that the effects of blurring from cross-subject misregistration will be applied twice (Coalson et al., 2018).

Unfortunately, no direct method for surface projection of a group average volume result is accurate. Once data has been averaged and smoothed in the volume, precise cortical localization cannot be recovered for cortical areas of high folding variability (i.e. association cortex) (Coalson et al., 2018) - although the magnitude of this problem may be minimal in some areas (like parts of the insula) where there is lower variability of folds and of areas relative to folds across subjects (Coalson et al., 2018). Therefore, the only way to accurately map the cortical areas for a statistical contrast of interest is following a surface-based preprocessing of individual subjects and surface-level group analysis. This fact is one of the major reasons we developed ciftify. This has implications for the appropriate combination of ciftify utilities when evaluating the result of a volume-based analysis: while it is technically possible to input a dense scalar file generated from ciftify_vol_result into ciftify_statclust_report, the resulting table should be interpreted with great caution. In particular, due to individual differences in cortical folding patterns and functional organization, the MMP1.0 parcellation cannot be accurately translated into MNI volume space (Coalson et al., 2018). Therefore, ciftify_peaktable is not recommended for the results from a volume-based analysis.

### 4.5 Future directions

While important advances in software development for the CIFTI file format have been achieved, a multitude of analysis workflows are more easily applied in volume than on the surface. Examples of comprehensive analysis packages include the NiLearn project for machine learning in Python, which contains several integrated utilities of NIFTI file manipulation, plotting, and time-series extraction (Abraham et al., 2014). Future work will be needed to integrate CIFTI file reading and writing into these programming environments. In addition, neuroinformatics platforms are well known for the systematic (machine-readable) reporting and meta-analysis of statistical maps in MNI (volumetric) space. Specific projects include neurosynth (neurosynth.org; Yarkoni et al., 2011), NeuroVault (https://neurovault.org/; Gorgolewski et al., 2015) and BrainMap (brainmap.org). Moreover, the greater neuroimaging community has aided these efforts by defining standards for reporting of these data as part of the neuroimaging data model (Maumet et al., 2016). The seeds for an alternative approach are provided by the BALSA database (balsa.wustl.edu, (Van Essen et al., 2017), a repository for sharing CIFTI-based results (via scene files in Connectome Workbench). Future work will be needed to develop methods for meta-analytic synthesis of these results.

Currently, the ciftify workflow employs surface anatomy (sulcal depth) based alignment to align cortical data across subjects. Recent advances show that improvements in cross-subject alignment can be gained using additional features such as resting-state connectivity (Conroy et al., 2013; Robinson et al., 2014; Tong et al., 2017) and/or surface myelin maps (Robinson et al., 2014). The HCP pipelines have implemented these advances in registration into their workflow, using the Multimodal Surface Matching registration tool (Robinson et al., 2018, MSM; 2014), to incorporate fMRI and anatomical features in the “MSMAll” registration (Glasser et al., 2016b). A ciftify workflow using only T1w data cannot match the precision of the HCP Pipeline surfaces. The HCP surface refinement procedure requires the high-resolution T2w image as well as additional tools for registration. Individualized parcellations based on functionally-related features (such as resting-state connectivity) could potentially provide an additional mechanism for improving cross-participant correspondence when the signal is averaged within these parcels (Glasser et al., 2016a; Wang et al., 2015). Future work is needed to validate which cortical regions might be effectively parcellated using the features available in legacy datasets.

There is rapid growth in the field of surface-based MR analysis. A growing number of analytic toolkits and methods use surface-based approaches, including novel 2D registration algorithms (Robinson et al., 2018; Tong et al., 2017), diffusion embedding(Margulies et al., 2016), and individualized parcellation(Glasser et al., 2016a; Kong et al., 2018; Schaefer et al., 2018; Wang et al., 2015)). However, due to differences in the underlying surface meshes (for example fsaverage5 vs fsLR32k vs CIVET), combining and comparing these methods remains technically challenging. The ciftify package allows for immediate application of tools and atlases developed using the HCP data (published in fsLR32k space) to a multitude of additional datasets. We believe the field will benefit from a simplified framework that promotes interoperability of different surface-based tools and facilitates evaluation of their performance on various type of MR acquistions (where diverse sources of noise may be present). We hope that the ciftify toolkit will provide a more standardized (and available) approach for the steps required to integrate fsLR32k atlases and tools into extended harmonized workflows.

### 4.6 Conclusions

The ciftify package offers a bridging solution for legacy data that will allow many researchers to adopt CIFTI format analyses. We intend for the analytic framework provided within the ciftify package to be a starting point for future bridging work. In parallel, the research community is increasingly embracing the HCP’s standards for MR data acquisition. Several initiatives, e.g. HCP Lifespan, the Connectomes Related to Human Disease, ABCD (Bjork et al., 2017) and the UK Biobank (Miller et al., 2016)) are now taking advantage of higher spatial and temporal resolution of MR data made possible using multi-band (multi-slice) acquisition (Ugurbil et al., 2013). Going forward, we hope that studies of such methodological quality will become the norm, rather than the exception. Whether using cifity for legacy data or CIFTI for HCP-quality data, it is our hope that an analytic shift from traditional volume-based approaches to surface-based (i.e. grayordinate) approaches will accelerate within the neuroimaging community.

## ACKNOWLEDGEMENTS

ANV acknowledges support related to the present work the Canadian Institute for Health Research, Ontario Mental Health Foundation, and Centre for Addiction and Mental Health Foundation, the Ontario Ministry of Research and Innovation, the Canadian Foundation for Innovation. MFG and DCVE acknowledge support from NIH (F30 MH097312 to MFG, RO1 MH-60974 to DCVE, and 3R24 MH108315 to DCVE).

---

**Supplementary Table 1:** the expected input directory structure (two options)

|  |  |
| --- | --- |
| The expected input structure (BIDS raw) |  |
| Description | Filename |
| T1w image | sub-{subject}/anat/sub-{subject}_T1w.nii.gz<br>sub-{subject}/anat/sub-{subject}_T1w.json |
| Functional image | sub-{subject}/func/sub-{subject}_task-{task}_bold.nii.gz<br>sub-{subject}/func/sub-{subject}_task-{task}_bold.json |
| Alternative Derivative input structure (if working from BIDS common derivatives)<br>*Note: is these file are not present in the /derivatives/ folder, then they will be generated by fMRIPrep* |  |
| Freesurfer Outputs | derivatives/freesurfer/sub-{subject}/ |
| Preprocessed T1w Image (i.e. the image the functional data is registered to) | derivatives/{func-pipeline}/sub-{subject}/anat/sub-{subject}_desc-preproc_T1w.nii.gz |
| Preprocessed functional image | derivatives/{func-pipeline}/sub-{subject}/func/sub-{subject}_task-{task}_space-T1w_desc-preproc_bold.nii.gz |

**Supplementary Table 2:**
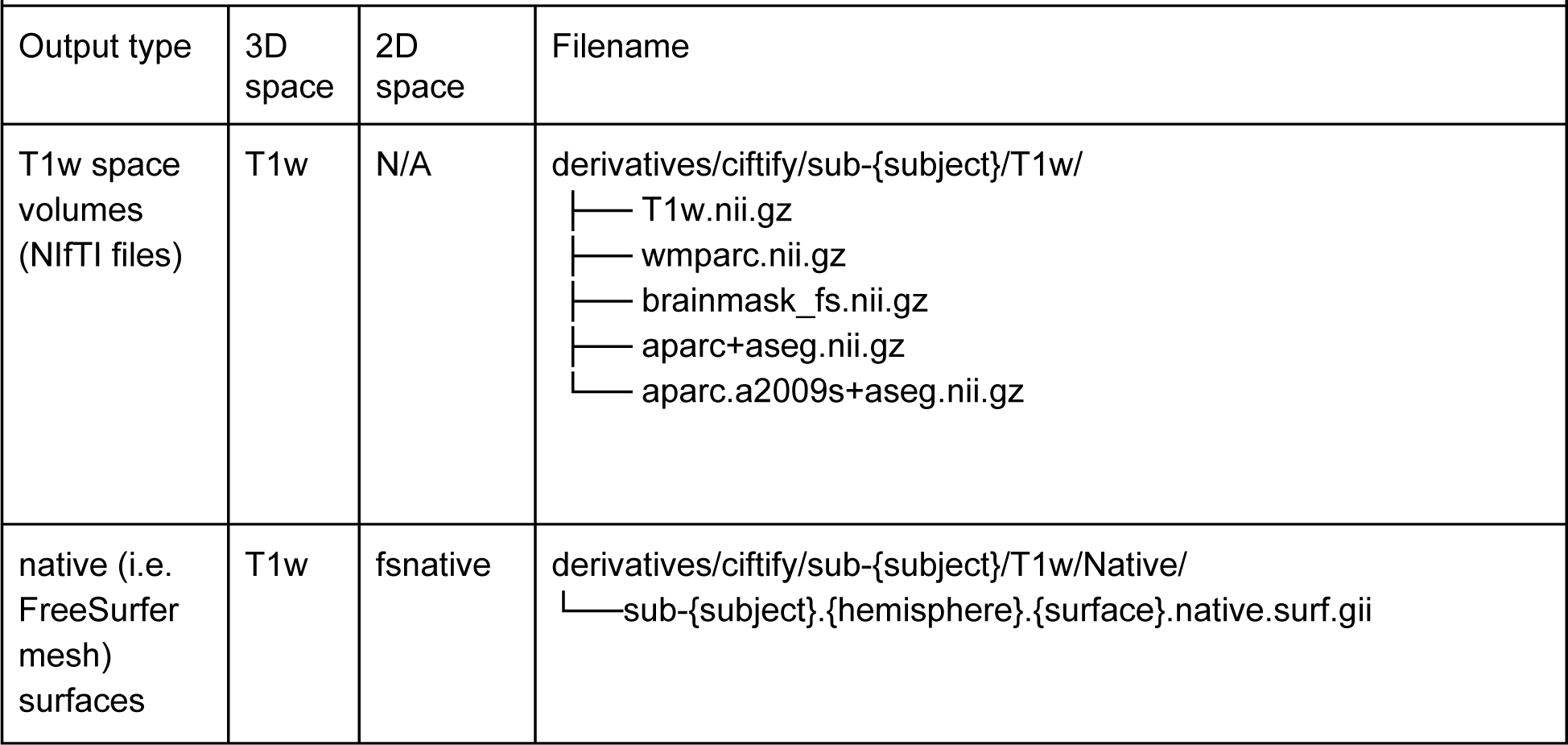
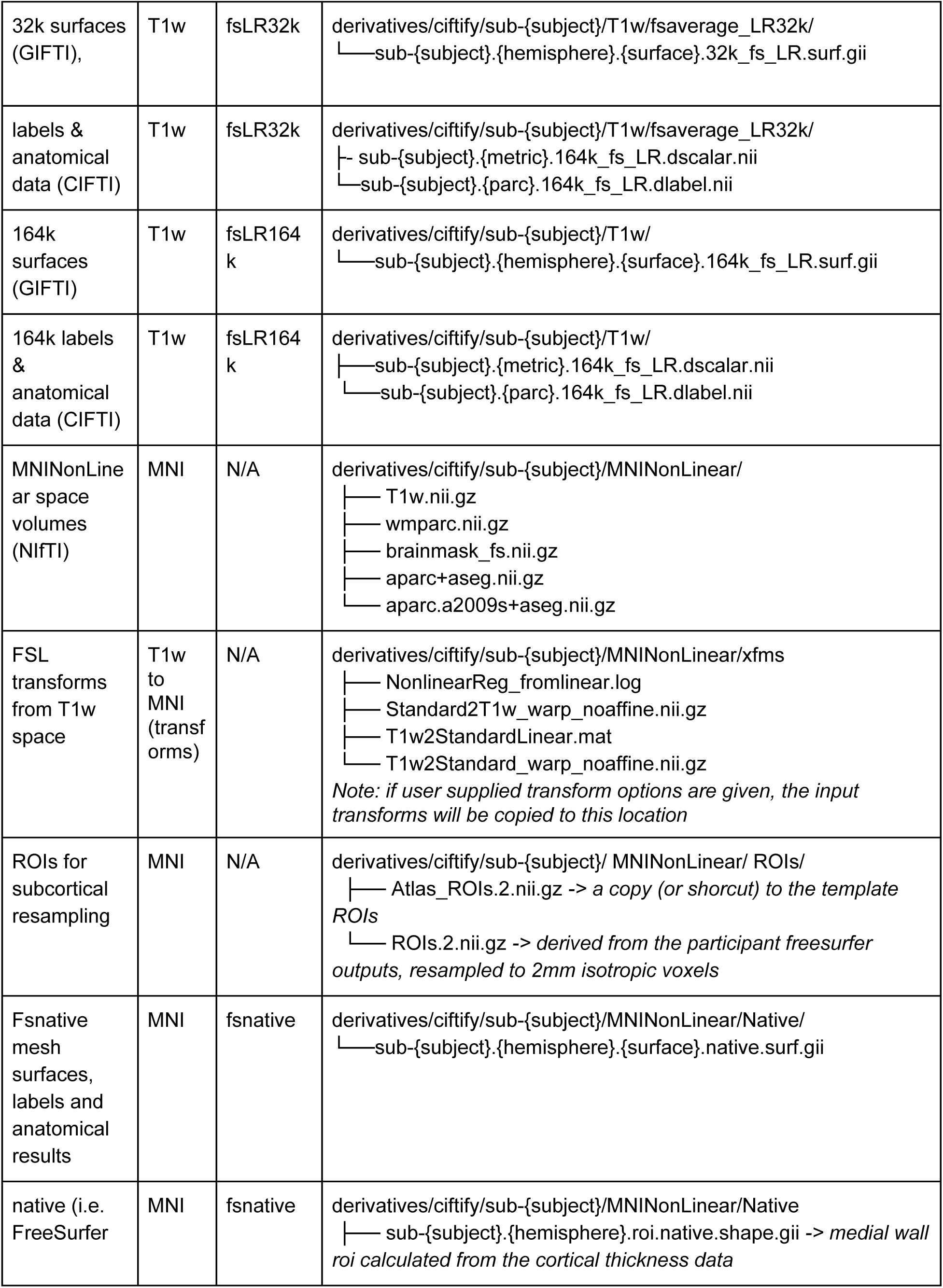
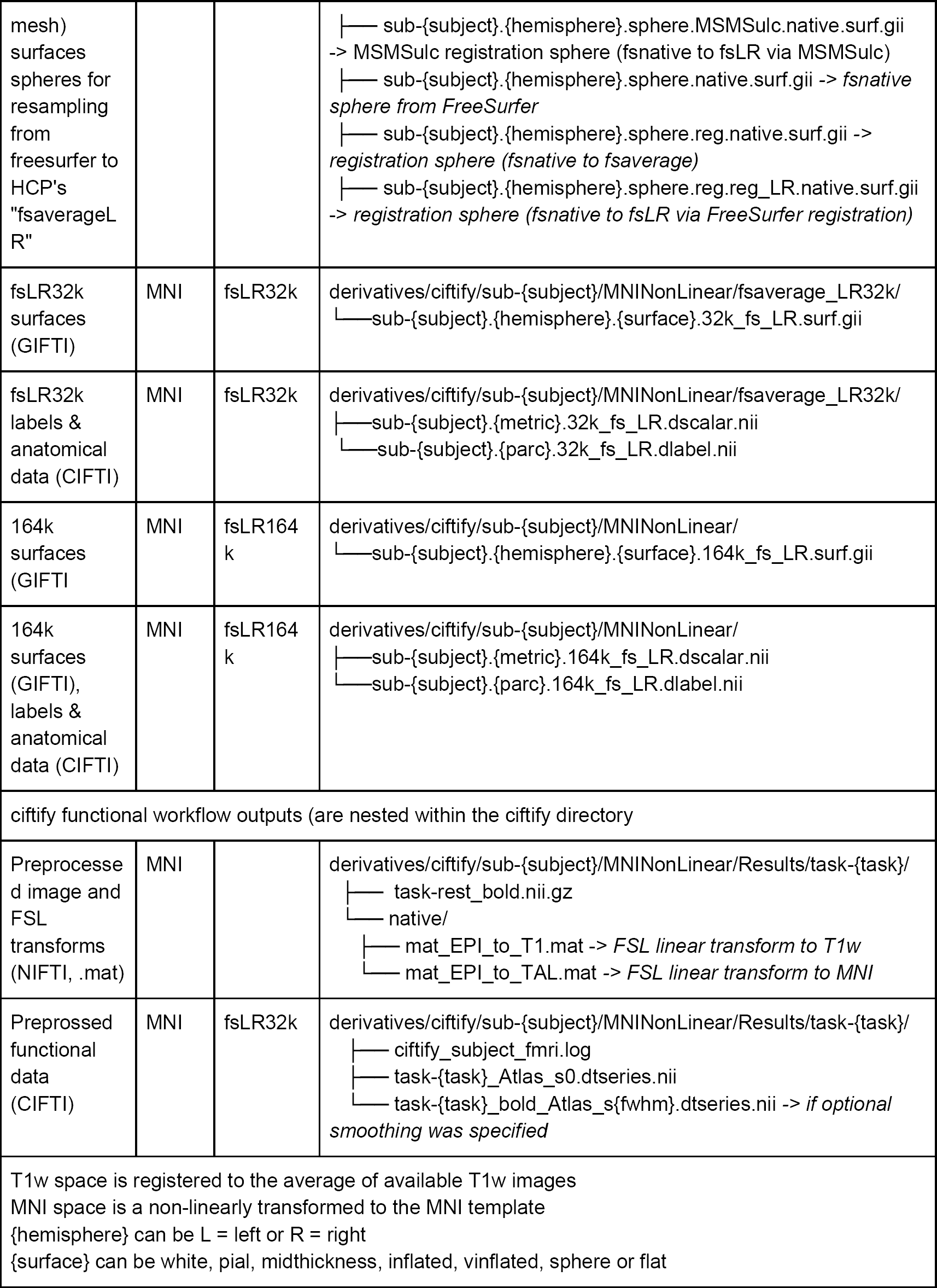
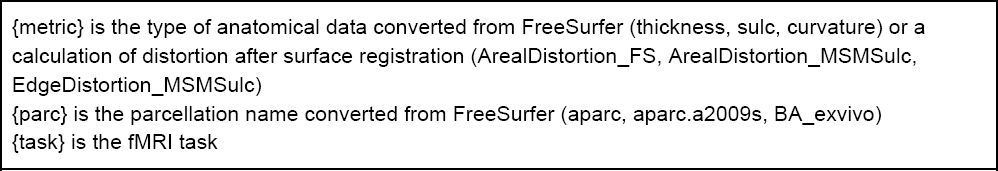
Details of the ciftify output directory structure *Similar to the output structure of the HCP-Pipelines*

